# The antipsychotic medications aripiprazole, brexpiprazole and cariprazine are off-target respiratory chain complex I inhibitors

**DOI:** 10.1101/2023.04.02.535223

**Authors:** Rachel E. Hardy, Injae Chung, Yizhou Yu, Samantha H.Y. Loh, Nobuhiro Morone, Clement Soleilhavoup, Marco Travaglio, Riccardo Serreli, Lia Panman, Kelvin Cain, Judy Hirst, Luis M. Martins, Marion MacFarlane, Kenneth R Pryde

## Abstract

Antipsychotic drugs are the mainstay of treatment for schizophrenia and provide adjunct therapies for other prevalent psychiatric conditions, including bipolar disorder and major depressive disorder. However, they also induce debilitating extrapyramidal syndromes (EPS), such as Parkinsonism, in a significant minority of patients. The majority of antipsychotic drugs function as dopamine receptor antagonists in the brain while the most recent ‘third’-generation, such as aripiprazole, act as partial agonists. Despite showing good clinical efficacy, these newer agents are still associated with EPS in ∼5-15% of patients. However, it is not fully understand how these movement disorders develop. Here, we combine clinically-relevant drug concentrations with mutliscale model systems to show that aripiprazole and its primary active metabolite induce mitochondrial toxicity inducing robust declines in cellular ATP and viability. Aripiprazole, brexpiprazole and cariprazine were shown to directly inhibit respiratory complex I through its ubiquinone-binding channel. Importantly, all three drugs induced mitochondrial toxicity in primary embryonic mouse neurons, with greater bioenergetic inhibition in ventral midbrain neurons than forebrain neurons. Finally, chronic feeding with aripiprazole resulted in structural damage to mitochondria in the brain and thoracic muscle of adult *Drosophila melanogaster* consistent with locomotor dysfunction. Taken together, we show that antipsychotic drugs acting as partial dopamine receptor agonists exhibit off-target mitochondrial liabilities targeting complex I.

## Introduction

Schizophrenia is a serious neurological disorder affecting 10 to 40 in 100,000 individuals (1). Pharmacological alleviation and prevention of psychosis is the primary clinical goal. Hyper-active dopamine signalling in the mesolimbic dopaminergic circuit is proposed to contribute to the pathology of schizophrenia (2). Therefore, antagonists of dopamine D2 and D3 receptors (D2R/D3R) represent a viable treatment approach (3). First- and second-generation D2R/D3R antagonists (e.g. haloperidol and risperidone) are therapeutically effective but also cause dose-limiting, serious neurological adverse events in a substantial proportion of patients (∼20-40%) (4). They include severe extrapyramidal symptoms (EPS) such as dystonia, akathisia, Parkinsonism, and tardive dyskinesia, which frequently lead to suspension of treatment (5, 6, 7–11). The side effects result from the on-target mechanism-of-action (MoA), which can suppress dopamine signalling to pathological levels and impair the nigrostriatal regulatory circuit that controls motor function. To reduce the incidence of neurological adverse events, third-generation antipsychotic drugs with a novel MoA have been developed. Specifically, aripiprazole (first approved for schizophrenia in the USA in 2002, with approximately 6 million prescriptions in 2017 (12)) functions as a partial D2R/D3R agonist and alleviates hyper-activation of dopaminergic circuits by competing for receptor binding with dopamine. It also prevents D2R/D3R hyper-activation during periods of excessive pre-synaptic dopamine release (as observed in the striatum of schizophrenic patients) (13, 14). Pharmacologically, the partial agonists succeed because their slow *k*_off_ rates mean that they remain bound to the D2R/D3R – even when dopamine levels transiently spike to pathological levels (15).

Aripiprazole has been approved for an expanded number of indications in both adult and juvenile patient populations: for bipolar disorders (suppression of manic episodes), major-depressive disorder and autism (16–19). Clinically, aripiprazole exhibits improved tolerability with a decreased frequency of neurological side effects (∼5-15%) (20–22). However, an unexpected minority of patients still experience severe movement disorders, both upon starting treatment and after prolonged treatment. In some instances, neurological dysfunction is irreversible following aripiprazole discontinuation (23, 24). The molecular mechanisms driving neurotoxicity remain unclear, but these clinical observations could suggest that aripiprazole exhibits an underlying off-target effect that drives permanent neuronal damage in a minority of patients.

Mitochondria are essential for ATP production in neurons and integral to their function and homeostasis (25). Mitochondrial dysfunction is broadly implicated in human neurodegeneration (26), affecting neurons that control movement. Specifically, mitochondrial activity is decreased in post-mortem brains of Parkinson’s disease (PD) patients (27–29), mutations in proteins controlling mitochondrial homeostasis causes early-onset PD (30, 31), and treating animals with the mitochondrial respiratory-chain complex I inhibitors rotenone or MPP^+^ recapitulates PD pathophysiology (32, 33). More broadly, mitochondrial inhibition is an important off-target mechanism of drug-induced-toxicity, contributing to drug attrition in clinical trials as well as drug withdrawal following approval due to severe adverse events (34, 35).

The interactions between aripiprazole and mitochondria are not understood. Here, we investigated whether off-target mitochondrial toxicity could be an underlying cause of adverse motor side-effects in patients taking aripiprazole. Using a suite of cellular and cell-free biochemical assays, we demonstrate that aripiprazole and its primary active metabolite, dehydroaripiprazole, are off-target cytotoxic mitochondrial respiratory-chain inhibitors. Both compounds selectively inhibited respiratory-chain complex I, at patient-relevant concentrations, and we defined inhibition of the complex I ubiquinone-binding site (Q-site) as the molecular-initiating-event (MIE) driving toxicity. Feeding *Drosophila melanogaster* food supplemented with the complex I inhibitor aripiprazole induced mitochondrial ultrastructural abnormalities in the brain and thoracic muscle that were linked with the development of significant locomotor defects. Finally, we reveal that two further clinically-approved third-generation antipsychotics, brexpiprazole and cariprazine, also specifically inhibit the Q-site of complex I. These data demonstrate that aripiprazole, brexpiprazole and cariprazine share a common off-target mechanism that drives mitochondrial and cellular toxicity.

## Results

### Aripiprazole and its primary active metabolite are toxic to cells in a manner consistent with mitochondrial toxicity

To determine the propensity of individual antipsychotic drugs for off-target mitochondrial toxicity, we employed the glucose-to-galactose metabolic shift assay (36). Briefly, galactose metabolism provides no net gain of ATP molecules from glycolysis, forcing the cell (particularly tumour cells that consume large amounts of glucose due to their Warburg phenotype) to rely on mitochondrial oxidative phosphorylation for ATP production. Thus, diverse mitochondrial toxicants can be uncovered using this model.

We treated human neuroblastoma cells (SH-SY5Y) cultured in glucose or galactose-containing media with 10 μM of each of 11 antipsychotic drugs from different classes for 18 h (Fig. 1A). Total cellular ATP levels were used as an endpoint to identify potential mitochondrial toxicants, with a selective reduction in galactose-conditioned cells as the criterion. Our screen showed that cellular ATP levels were selectively reduced in galactose-conditioned cells only by aripiprazole (Fig. 1A). Although chlorpromazine also induced a modest decline in cellular ATP levels, this was observed in both glucose and galactose-cultured cells. No effects were observed with iloperidone, lurasidone, haloperidol, risperidone, olanzapine, quetiapine, clozapine, zotepine or paliperidone. 2-Deoxyglucose (2-DG; a well-characterized inhibitor of glycolysis), piericidin A (a potent complex I inhibitor) and antimycin A (a potent complex III inhibitor) were used as positive controls.

**Figure 1.**
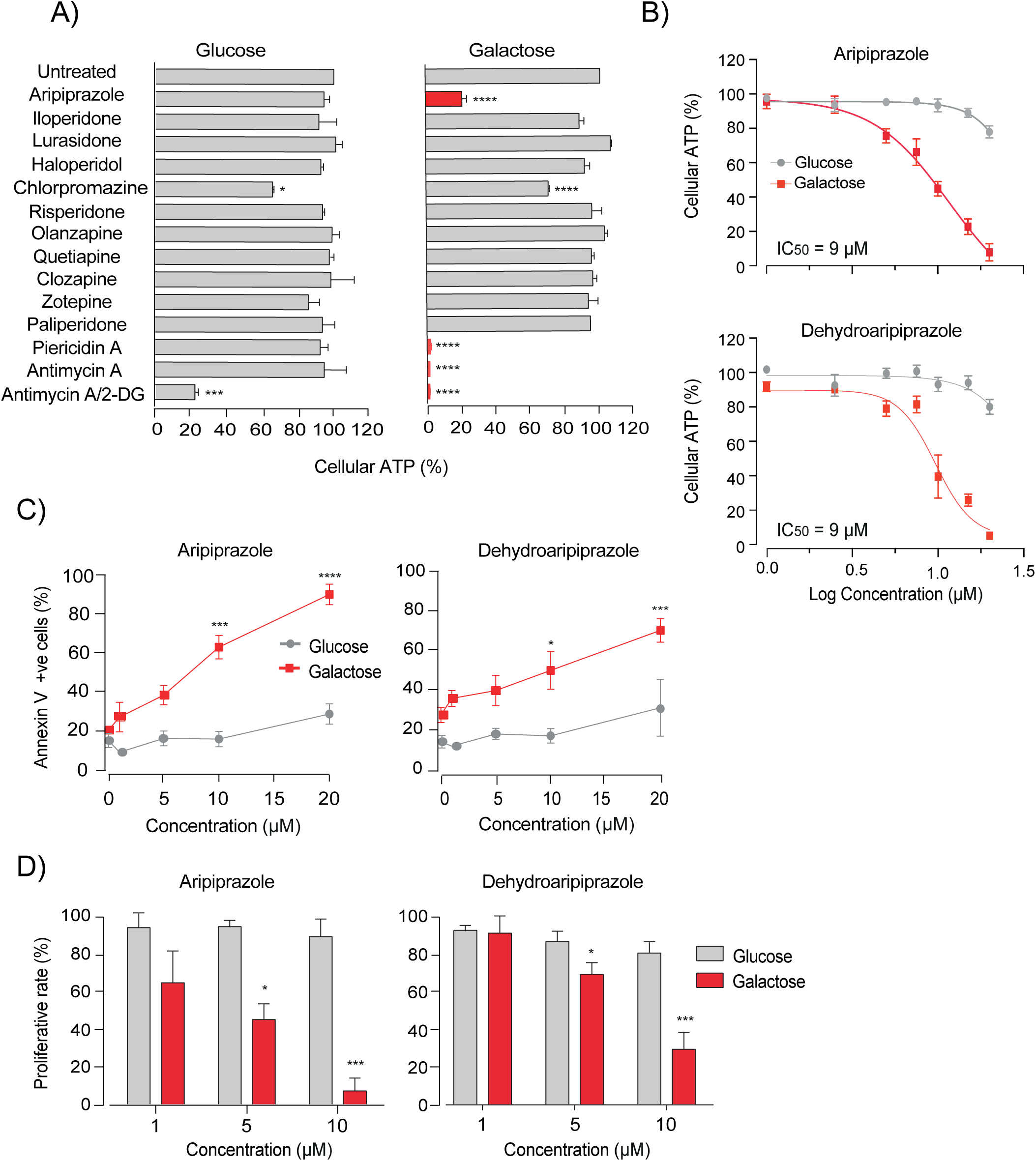
Aripiprazole and its primary active metabolite are selectively toxic to galactose-conditioned SH-SY5Y cells. (A) Normalised cellular ATP measurements from glucose- and galactose-conditioned SH-SY5Y cells exposed to the indicated antipsychotic drugs (10 μM), piericidin A, antimycin A (5 µM) or 2-deoxyglucose (2-DG) (50 mM) for 18 h (the data are mean averages ± range from 2 independent experiments with 3 technical repeats; asterisks show the results from one-way ANOVA with Dunnett’s multiple comparison test, normalised to control). (B) Normalised cellular ATP measurements from glucose- and galactose-conditioned SH-SY5Y cells treated with the indicated concentrations of aripiprazole or dehydroaripiprazole for 18 h. Data were fitted to the standard dose-effect relationship (activity (%) = = Bottom + (Top-Bottom)/(1+10^((LogIC50-X)*HillSlope)) using GraphPad Prism version 8.0 (mean ± SEM from 3 independent experiments, normalised to control). (C) Quantification of cell death in glucose- and galactose-conditioned SH-SY5Y cells treated with increasing concentrations of aripiprazole and dehydroaripiprazole for 18 h (mean ± SEM from 3 independent experiments, asterisks, one-way ANOVA with Dunnett’s multiple comparison test, normalised to control) (corresponds to Supp Figure 1). (D) Quantification of glucose and galactose-conditioned SH-SY5Y cell proliferative rates following treatment with indicated concentrations of aripiprazole or dehydroaripiprazole over an 86 h period. Proliferation rates were calculated from the linear phase of each growth curve (mean ± SEM from 3 independent experiments, asterisks, one-way ANOVA with Dunnett’s multiple comparison test, normalised to control) (corresponds to Supp Figure 2).

Next, concentration-titrations were performed with aripiprazole and its primary active metabolite dehydroaripiprazole. Levels of dehydroaripiprazole in patient serum are ∼40% of the parent compound (37). Therefore, it was important to clarify whether the galactose-selective toxicity of aripiprazole was retained following metabolism. Both compounds induced dose-dependent declines in cellular ATP levels in galactose-conditioned SH-SY5Y cells after 18 h treatment (Fig. 1B). The IC_50_ values (concentrations required for 50% inhibition) for aripiprazole and dehydroaripiprazole were both 9 μM, with concentrations to 5 μM inducing modest declines (20-30%) in cellular ATP levels.

Further investigation demonstrated that both aripiprazole and dehydroaripiprazole promote cell death in a galactose-selective manner. Both compounds (5 μM) induced cell death after 18 h treatment, with a significant increase at 10 μM (∼43% for aripiprazole and ∼20% for dehydroaripiprazole) (Fig. 1C and Supp. Fig 1). Furthermore, clear reductions in cellular proliferation rate were observed in galactose-conditioned SH-SY5Y cells treated with 5 or 10 μM of each compound over an 86 h period (Fig. 1D and Supp.2). Therefore, we have identified aripiprazole and its primary pharmacologically active metabolite to be selectively toxic to SH-SY5Y cells cultured in galactose-medium: they diminish cellular ATP, viability and proliferative rate in a manner consistent with mitochondrial toxicity.

### Aripiprazole and structurally related antipsychotic drugs inhibit ubiquinone reduction by mitochondrial respiratory complex I

Next, we sought to determine the direct effects of aripiprazole and dehydroaripiprazole on mitochondrial bioenergetic function. Based on profound similarities in chemical substructure, we also investigated the more recently approved antipsychotic compounds brexpiprazole and cariprazine, as well as didesmethylcariprazine: a primary active metabolite of cariprazine (16).

Extracellular flux analysis was employed to assess the effects of the drugs on mitochondrial respiration. The assays were carried out in serum-free media to enable the effects to be assessed without any interference from drug-serum binding; aripiprazole and dehydroaripiprazole are known to be highly bound (>99%) to plasma proteins in patients (38). In experiments using foetal bovine serum to compare SH-SY5Y cells cultured in serum-supplemented or serum-free galactose-media, 10 µM aripiprazole or dehydroaripiprazole both induced greater declines in ATP levels in cells in serum-free media (Supp. 3A), consistent with this fact.

The oxygen consumption rate (OCR), which depends on the function of the electron-transport chain (ETC), was measured in SH-SY5Y cells treated with varying concentrations of each drug (see Supp 3B for schematic of a standard mitochondrial respiration stress test). Importantly, we observed a significant decline in both basal and maximal OCR following a 4 h treatment with aripiprazole, dehydroaripiprazole, brexpiprazole or cariprazine, indicating that these drugs are direct mitochondrial toxins that inhibit the ETC (Fig. 2A and Supp. 3C). No change in OCR was observed with didesmethylcariprazine, even at the highest concentration of 10 μM.

**Figure 2.**
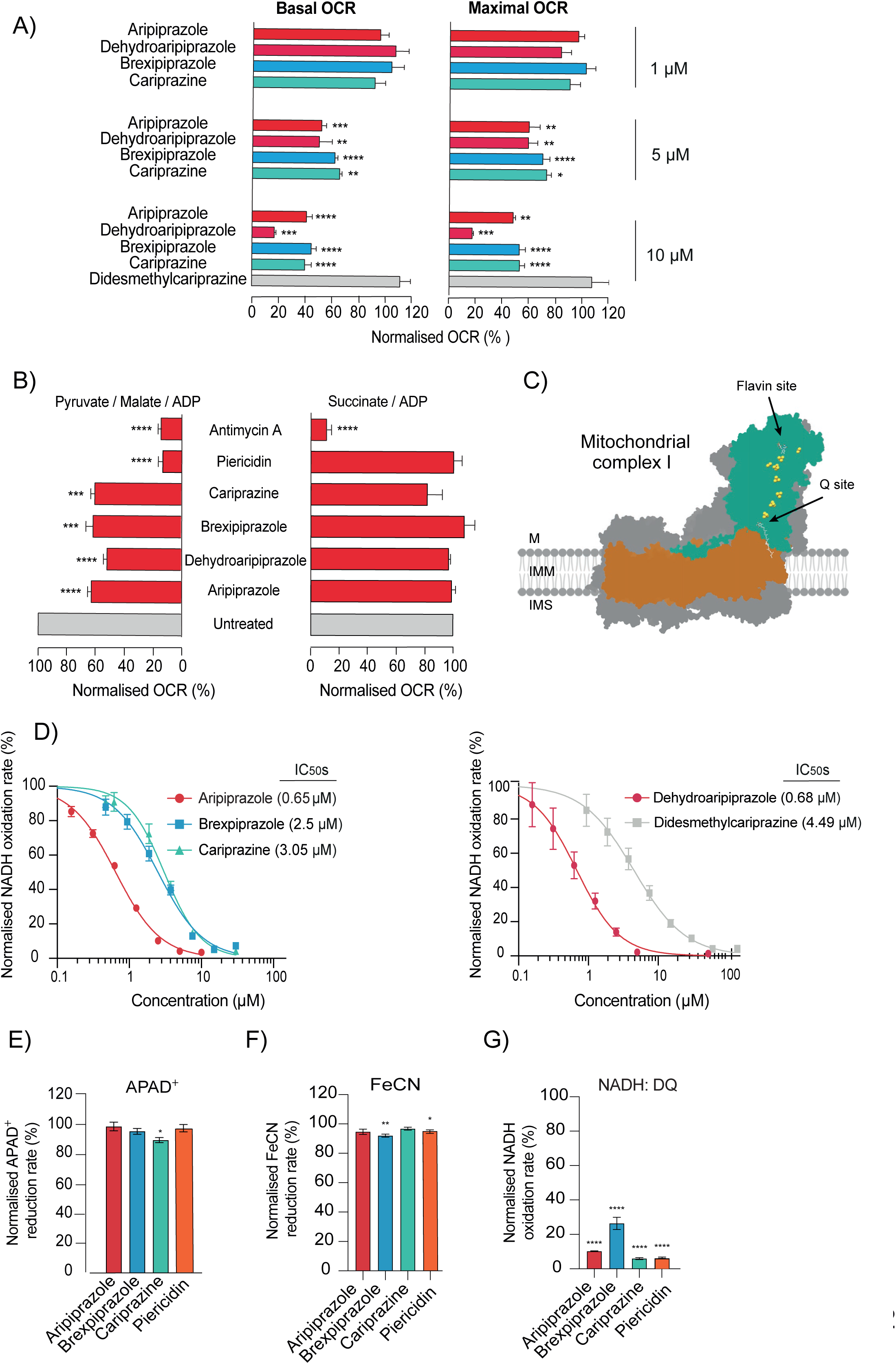
Aripiprazole, Brexpiprazole, and Cariprazine directly inhibit respiratory complex I *via* the Q-channel. (A) Normalised basal and maximal OCR measurements from SH-SY5Y cells treated for 4 h with indicated concentrations of aripiprazole, dehydroaripiprazole, brexpiprazole, cariprazine or didesmethylcariprazine (mean ± SEM from 4 independent experiments, asterisks, one-way ANOVA with Dunnett’s multiple comparison test, normalised to control) (corresponds to Supp. Figure 3C). (B) Normalised OCR measurements driven by complex I substrates (pyruvate/malate) or the complex II substrate (succinate). Permeabilised SH-SY5Y cells were treated with 10 μM aripiprazole, dehydroaripiprazole, brexpiprazole, cariprazine or 5 μM piericidin A, antimycin A (mean ± SEM from 4 independent experiments, asterisks, one-way ANOVA with Dunnett’s multiple comparison test, normalised to control) (corresponds to Supp. Figure 4B). (C) A cartoon depiction of mammalian complex I structure with the two substrate-binding sites indicated. (D) Normalised NADH:O_2_ oxidoreduction rates of bovine mitochondrial membranes exposed to varying concentrations of aripiprazole, brexpiprazole, cariprazine, dehydroaripiprazole and didesmethylcariprazine. Respective IC_50_ values from the standard dose-effect relationship (see Methods) are indicated in brackets. (E) Normalised NADH:APAD^+^, (F) NADH:FeCN, and (G) NADH:DQ oxidoreduction rates of isolated bovine complex I exposed to 100 μM aripiprazole, brexpiprazole, cariprazine or 1 µM piericidin A (mean ± SEM from 3 independent wells per treatment, asterisks, one-way ANOVA with Dunnett’s multiple comparison test, normalised to control).

Next, we sought to dissociate the toxic off-target effects of the drugs on mitochondria from their on-target effects on the dopamine D2R/D3R. Although these antipsychotics exert functionality at multiple serotonin receptor subtypes, we focused on dopamine receptors since these are the primary therapeutic target linked to the onset of EPS in patients (3, 39). First, SH-SY5Y cells were treated with an excess concentration of the D2R/D3R agonist dopamine or antagonist haloperidol. Neither of these chemicals induced a decline in basal or maximal OCR (Supp. 3D). In contrast, a 30 min pre-treatment with haloperidol or dopamine followed by a 30 min treatment with aripiprazole induced a significant decline in basal and maximal OCR. The retention of mitochondrial toxicity under conditions in which the D2R/D3R are expected to be fully occupied confirms that this is an off-target effect. Furthermore, aripiprazole was also shown to induce significant declines in basal and maximal OCR in human cardiomyocytes (HCMs), despite the fact that these cells express no detectable level of D2R and extremely low levels of D3R (Supp. 3E and 3F). We conclude that the drug-induced mitochondrial toxicity observed is independent of D2R/D3R partial agonism.

Next, to determine the molecular mechanism of antipsychotic-induced mitochondrial toxicity, we stimulated plasma-membrane-permeabilised cells with pyruvate and malate to induce complex I-linked respiration, and with succinate to stimulate complex II-linked respiration (see Supp. 4A for schematic representation of the ETC). Crucially, we observed a significant decrease in pyruvate/malate-driven respiration following treatment with aripiprazole, dehydroaripiprazole, brexpiprazole and cariprazine (10 μM) (Fig. 2B and Supp. 4B). Notably, a pre-treatment period of 2 h was required for inhibition by cariprazine to be observed. In contrast, no significant change in succinate-driven respiration was detected with any drug. The same pattern was observed using piericidin A, a specific complex I inhibitor (see Fig. 2C for a schematic of respiratory complex I and the major catalytic sites). To confirm the specificity for complex I, NADH and succinate oxidation rates were measured in bovine mitochondrial membranes. All three antipsychotic drugs, and both primary active metabolites, induced dose-dependent declines in NADH oxidation rate (see Fig. 2D), but did not affect succinate oxidation substantially (Supp. 4C). Although a significant decrease was observed with cariprazine and dehydroaripiprazole, this was at an excess concentration of 50 μM. Further analyses in bovine sub-mitochondrial particles confirmed that, as expected from its inhibition of the ETC, aripiprazole significantly inhibits ATP synthesis in a dose-dependent manner (Supp. 4D).

Next, we sought to determine the mechanism of antipsychotic-induced complex I inhibition. Complex I harbours two substrate-binding sites that are both candidate inhibitor binding sites: the NADH-binding site that contains a flavin mononucleotide cofactor (flavin-site) and the ubiquinone-binding site (Q-site) (Fig. 2C). To assess NADH-stimulated flavin-site activity, we measured the reduction rate of the artificial electron acceptors APAD^+^ and FeCN by the reduced flavin in isolated bovine complex I. We identified no substantial change in the NADH: APAD^+^ /FeCN oxidoreduction rates following exposure to 100 µM aripiprazole, brexpiprazole or cariprazine, clearly demonstrating that the flavin-site is not inhibited (Fig 2E and 2F). The artificial ubiquinone analogue decylubiquinone (DQ) was used assess the Q-site activity in isolated bovine complex I and these data confirmed that each drug induced a substantial decline in NADH:DQ oxidoreduction (Fig. 2G). Therefore, our findings support that dopamine receptor partial agonist antipsychotic drugs are toxic to mitochondria *via* inhibition of the Q-channel of respiratory complex I.

### Third-generation antipsychotics display off-target mitochondrial toxicity in primary mouse neurons

The antipsychotic drugs examined here are designed to modulate dopaminergic neurotransmission in the mesolimbocortical pathways of the human brain. To examine whether the off-target mitochondrial toxicity observed in human neuroblastoma cells is also observed in post-mitotic neurons, primary cultures of forebrain and ventral midbrain neurons were established from wild-type mouse embryos (E13.5). Isolation of these distinctive populations enabled us to compare the propensity for off-target mitochondrial toxicity in dopaminergic versus non-dopaminergic neuronal subtypes.

To confirm the dopaminergic phenotype of ventral midbrain neurons, cells were stained for the specific dopaminergic marker tyrosine hydroxylase (TH). As expected, ventral midbrain neurons stained strongly for TH, whilst forebrain neurons did not express detectable TH (Fig. 3A). Both neuronal subtypes were also stained with β3 tubulin and Hoechst after 7 d in culture, to confirm their viability (Fig. 3B). Cellular ATP levels were then measured in both neuronal subtypes, with and without treatment with each antipsychotic drug for 18 h. A significant decline in cellular ATP (approximately 20%) was observed in ventral midbrain neurons treated with aripiprazole, brexpiprazole and cariprazine (Fig. 3C). In contrast, no change in cellular ATP levels was recorded in forebrain neurons. Treatment with piericidin A significantly reduced ATP production in both midbrain and forebrain neurons (Fig. 3C).

**Figure 3.**
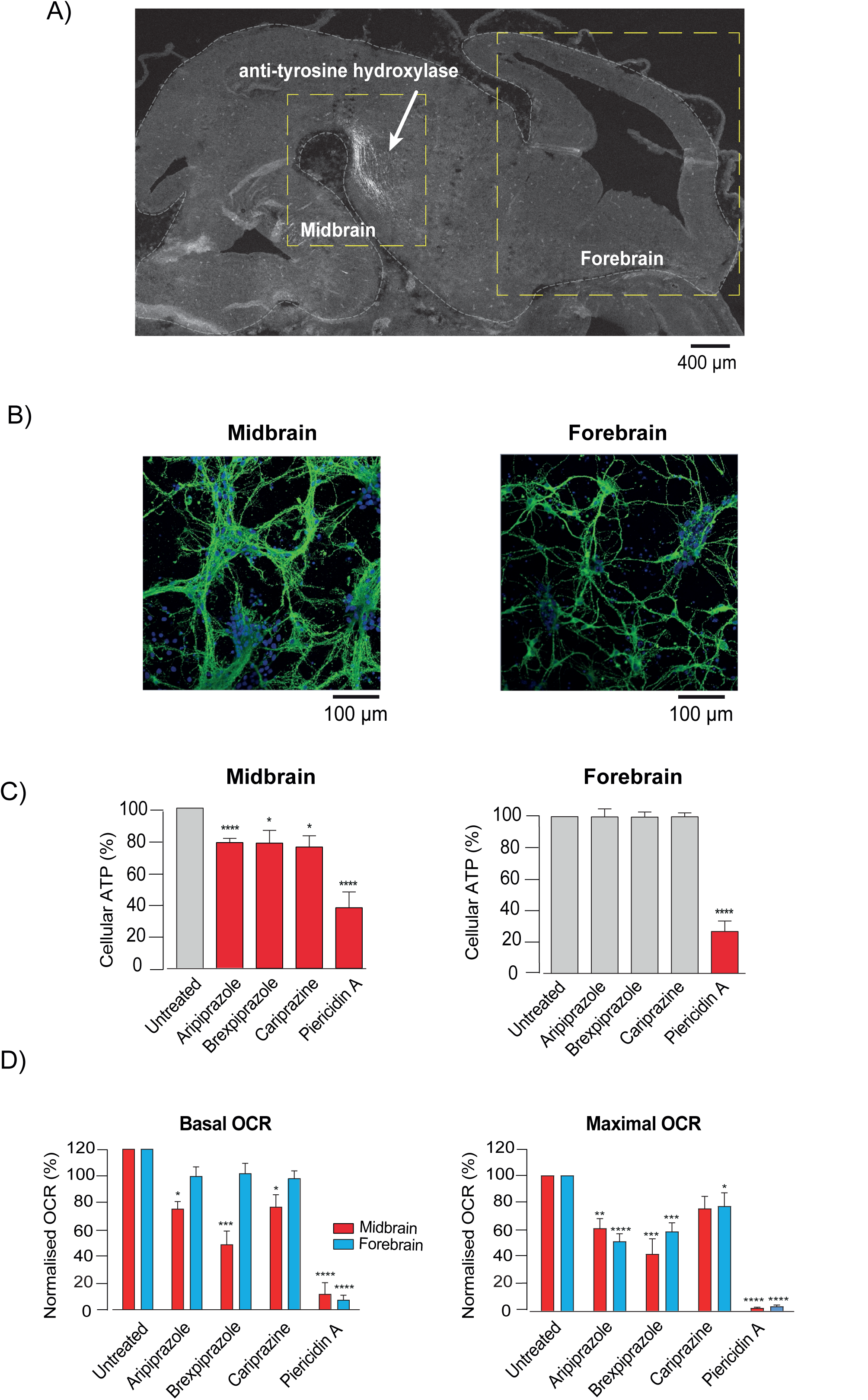
Aripiprazole, brexpiprazole and cariprazine are toxic to primary neonatal mouse neurons. (A) Representative image of a sagittal section from an embryonic mouse brain (e13.5) stained with an anti-tyrosine hydroxylase (TH) antibody to highlight dopaminergic neurons (white arrow). Dashed yellow boxes represent the ventral midbrain and forebrain areas used to prepare neurons. (B) Representative confocal images of ventral midbrain and forebrain neurons (e13.5) maintained for 7 d and stained with Hoechst (blue) and an antibody to β3 tubulin (green). (C) Normalised cellular ATP measurements from mouse embryonic (e13.5) ventral midbrain and forebrain neurons. Neurons cultured for 7 d were exposed to aripiprazole, brexpiprazole, cariprazine or piericidin A (5 μM for 18 h) (mean ± SEM from 5 independent experiments, asterisks, one-way ANOVA with Dunnett’s multiple comparison test, normalised to control). (D) Normalised basal and maximal OCR measurements from mouse ventral midbrain and forebrain neurons (e13.5) grown for 7 d and treated with aripiprazole, brexpiprazole, cariprazine or piericidin A (5 μM for 4 h) (corresponds to Supp Figure 5B) (mean ± SEM from 5 independent experiments, asterisks, one-way ANOVA with Dunnett’s multiple comparison test, normalised to control).

Importantly, analyses of NADH oxidation rate in mouse mitochondrial membranes demonstrated that all drugs of interest retained their inhibitory effects on complex I-linked respiration (Supp. 5A), confirming that mitochondrial toxicity is conserved across species. Next, we measured the direct effects of each antipsychotic drug on mitochondrial respiratory function. Crucially, each compound induced a significant decline in basal and maximal OCR in midbrain neurons (Fig. 3D and Supp. 5B). In contrast, while no statistically significant change in basal OCR was detected with either antipsychotic drug in forebrain neurons, a substantial and significant decline in maximal OCR was still recorded (∼28% aripiprazole, ∼42% brexpiprazole, ∼26% cariprazine). The fact that aripiprazole, brexpiprazole and cariprazine had different effects on the two cell types indicates differential rates of uptake and accumulation – perhaps increased in the ventral midbrain neurons resulting from their unmyelinated nature (40).

### Aripiprazole induces mitochondrial toxicity in *Drosophila melanogaster*

To explore the mitochondrial liabilities of third-generation antipsychotics in a living organism, we employed *Drosophila melanogaster* as an *in vivo* model. *Drosophila* also have the advantage that, effects on dopamine signalling can be distinguished from effects on mitochondrial function. To confirm that mitochondrial toxicity was conserved in flies*, Drosophila* S2 cells and mitochondria isolated from *Drosophila* were treated with aripiprazole, brexpiprazole and cariprazine for 4 h and 15 min, respectively. All drugs induced a significant decline in basal OCR (20-34%) and maximal OCR (27%- 74%) in S2 cells (Supp. 6A and 6B). Acute drug treatment also decreased basal OCR (and in the case of aripiprazole and brexipiprazole maximal OCR) of isolated mitochondria from *Drosophila*, while the canonical complex I inhibitor piericidin A completely abolished mitochondrial respiration (Supp. 6C and 6D). Taken together, these data demonstrate that third-generation antipsychotic drugs induce mitochondrial toxicity in *Drosophila melanogaster*, demonstrating the utility of this model for our study.

Due to limitations in compound availability, aripiprazole was the only drug investigated in behavioural analyses. First, it was important to distinguish the on-target versus off-target effects of aripiprazole in *Drosophila.* That is, behavioural/functional effects caused by the partial agonist interaction of aripiprazole with its primary therapeutic target (D2R/D3R) versus interaction with an unintended molecular target (e.g. mitochondrial complex I). In *Drosophila,* dopamine acts as an arousal signal by downregulating the activity of the dorsal layer of fan-shaped body (dFB) neurons, an effector component of the sleep homeostat (41). Sleep fragmentation correlates with increased dopamine signalling in flies and can be measured by monitoring the activity of individual flies (42). Therefore, we used sleep fragmentation as a way to measure the on-target effect of the drug. Accordingly, significant increases in the number of inactivity episodes were detected in aripiprazole-fed flies after 14 and 21 d of exposure (Fig. 4A).

**Figure 4.**
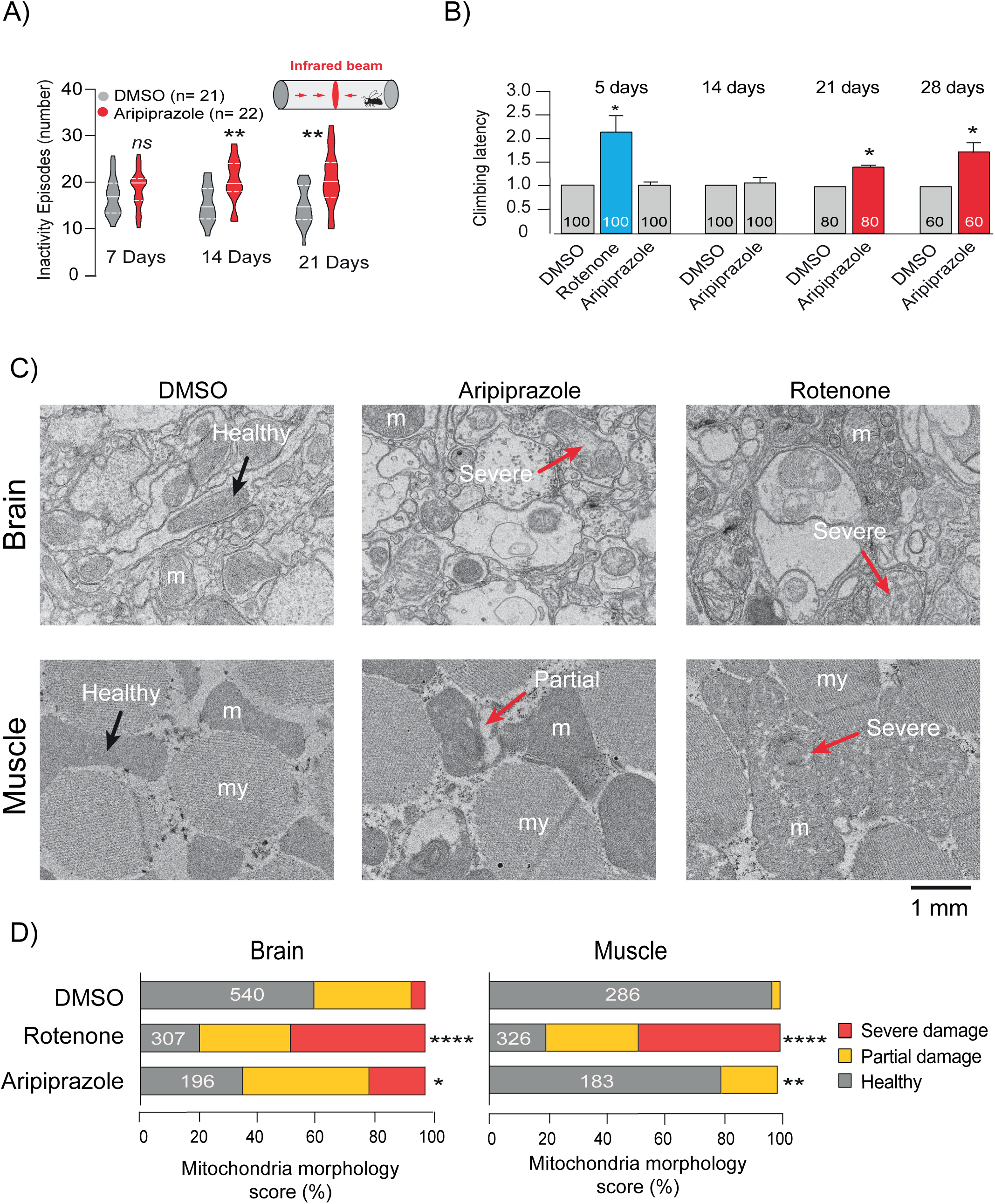
Aripiprazole causes motor dysfunction and mitochondrial defects in adult flies. (A) Total activity of adult flies treated with DMSO (0.5% v/v) or aripiprazole (1 mM). Measured for indicated periods using a Trikinetics system (*n* = number of flies per test group, asterisks, two-way ANOVA with the Tukey multiple comparison test). (B) Climbing performance of adult flies fed with DMSO (0.5% v/v) rotenone or aripiprazole (1 mM) for indicated periods. Total number of flies tested per group are indicated on the graph (mean ± SEM from 3 independent experiments, asterisks, two-tailed unpaired *t*-test). (C) Representative TEM images of brain and thoracic muscle from adult flies treated with DMSO (0.5%) or aripiprazole (1 mM) for 28 d or rotenone (1 mM) for 5 d (arrows, examples of the scoring criteria for the morphological assessment, m = mitochondria, my = myofibrils). (D) Quantification of mitochondrial damage. The total number of mitochondria scored in each sample group is indicated (samples pooled from 2 flies per treatment, chi-squared analysis with Bonferroni correction).

Mitochondrial dysfunction in the skeletal muscle has been shown to impair motor performance in *Drosophila* (43). To assess the effect of mitochondrial complex I inhibition on this phenotype, we measured the climbing ability of adult flies fed with aripiprazole or the canonical complex I inhibitor rotenone for varying amounts of time: with climbing latency defined as the amount of time taken for 5 flies (out of 20) to reach a 15 cm height marker within a vertical chamber. Therefore, an increase in climbing latency reflects impaired locomotion. Although no climbing impairment was noted in aripiprazole-fed flies after 5 or 14 d feeding, a significant increase in climbing latency was detected after 21 and 28 d (Fig. 4B). A robust and significant increase in climbing latency was also detected in flies exposed to rotenone for 5 d (later time-points are not shown, as flies did not survive the treatment). Finally, we investigated the effect of aripiprazole treatment on the ultrastructure of mitochondria in the brain and thoracic muscle of adult flies. Various studies have shown DMSO to be toxic to flies, with 0.5% DMSO only appearing to induce damage in our study following chronic feeding (44). Although rotenone induced clear perturbations in mitochondrial ultrastructure, with abundant cristae fragmentation present in both brain and thoracic muscle after 5 d feeding, no significant damage was detected with aripiprazole (Supp. 6E). In comparison, clear abnormalities in mitochondrial ultrastructure were evident in the brain and thoracic muscle of aripiprazole-fed flies after 28 d (Fig. 4C-D). Clear areas of vacuolisation were detected in ∼20% of densely packed mitochondria in thoracic muscle, with no concurrent damage to myofibrils being noted. Similarly, a significant number of brain mitochondria (∼20%) presented with fragmented cristae following 28 d aripiprazole feeding, with a mixture of partial and severe damage being observed (Fig. 4C-D). It should be noted that ∼40% of mitochondria appeared damaged in the brains of flies fed DMSO for 28 d, and the amount of damage induced by aripiprazole was calculated relative to this value. Taken together, our data show that *in vivo* exposure to aripiprazole results in mitochondrial ultrastructural damage in both the brain and thoracic muscle of adult flies, which is linked with impaired motor performance.

## Discussion

First-generation antipsychotic drugs are potent D2R antagonists inducing severe and high-frequency EPS (∼30-70% of all patients) (10, 11, 45). To improve tolerability and lessen patient suffering, the D2R/D3R partial agonists aripiprazole, brexpiprazole and cariprazine were developed, with this novel MoA promoting favourable drug side-effect profiles. However, adults taking these drugs still exhibit a high incidence (∼5-15%) of EPS in both clinical trials and real-world analyses (16, 21, 46–48). EPS were similarly high (∼15-25%) in the clinical trials of paediatric patients (10 to 17 years) treated with oral aripiprazole for bipolar mania, demonstrating that neurological toxicity is conserved from children to adults (21). Moreover, EPS are irreversible in some adults and children after discontinuation of third-generation antipsychotic drugs, suggesting these medications comprise toxic properties that can inflict permanent neuronal damage (23, 24). Given that dopamine receptor partial agonists are prescribed to millions of patients globally each year, it is crucial to better understand why EPS remain a considerable liability for these medications and to delineate if they are driven by on- or off-target pharmacology. Here, we investigated if aripiprazole, brexpiprazole and cariprazine affect mitochondrial function (respiration), and explored the possible contribution of this effect to human EPS.

Our data show that aripiprazole, brexpiprazole and cariprazine are direct inhibitors of the ETC in cells (SH-SY5Y cells and primary mouse neurons) and mitochondrial membranes (bovine and mouse). In a cellular context, none of the other antipsychotic medications screened here induced mitochondrial toxicity. Whilst our findings are contrary to previous reports that detected inhibition of the ETC by a wider range of antipsychotic compounds, the earlier studies were limited to cell-free purified mitochondrial systems (49, 50). We propose that first and second-generation antipsychotic compounds cannot gain access to their molecular target(s) in mitochondria in intact cellular systems, as tested here. This underlines the importance of combining mitochondrial toxicity data in cell-free and intact cellular assays (49–53). Using functional assays on respiratory complexes, we show the three partial agonists specifically inhibit the site of ubiquinone reduction (Q-site) in mitochondrial complex I. We present strong evidence to show that this mitochondrial inhibition is off-target and disconnected from the on-target pharmacology of third-generation antipsychotic drugs. All three compounds inhibit complex I activity in isolated complex I, cell-free purified mitochondrial membranes and intact cellular systems. In contrast, respiration in cells is not inhibited following treatment with dopamine (full agonist of D2R and D3R) or haloperidol (D2R and D3R antagonist). Importantly, aripiprazole, brexpiprazole and cariprazine inhibit mitochondrial respiration in both primary neurons and non-neuronal cells (human cardiomyocytes) that lack D2R expression.

*In vivo* studies using *Drosophila melanogaster* showed that chronic feeding with aripiprazole resulted in significant climbing defects after 21 d, which closely paralleled the emergence of severely damaged mitochondria in the thoracic muscle and brain. These defects further deteriorated during 21-28 d treatment, suggesting that aripiprazole elicits degenerative toxicity. Importantly, the delayed onset of locomotor dysfunction in flies mirrors aripiprazole clinical data, with EPS typically presenting weeks-to-months after initiating aripiprazole treatment. The canonical complex I inhibitor rotenone also induced mitochondrial cristae fragmentation and locomotor dysfunction, albeit with significantly faster onset. These differential rates of toxicity between rotenone and aripiprazole are expected, because rotenone more potently inhibits complex I compared to aripiprazole (IC50 value of 6.9 nM using bovine mitochondrial membranes, more than 100x lower than aripiprazole (54)). Crucially, we detected our on-target D2R-dependent marker (i.e. increased inactivity episodes) at an earlier time-point (14 d) compared to the climbing defects (21 d), confirming that mitochondrial toxicity is disconnected from D2R/D23R partial agonism in a living organism.

Although further studies are required to unequivocally show that motor side-effects are a direct consequence of mitochondrial inhibition, further evidence exists to support this proposal. Aripiprazole occupies approximately 95% of D2Rs/D3Rs in the human striatum (≥14 d treatment), with extremely strong binding affinities (*K*_i_ = 0.34 nM for D2R and 0.8 nM for D3R, among the highest recorded for any antipsychotic drug (55, 56)). A lower dopamine receptor occupancy (75-80%) by first-generation antipsychotics (i.e. D2R/D3R antagonists) promotes serious EPS (57–59). Therefore, if aripiprazole does induce motor disorders by D2R/D3R partial agonism, it is surprising that the prevalence is relatively low. This alone suggests that an alternative off-target mechanism is responsible for driving aripiprazole-induced neurotoxicity and associated EPS.

We next considered if the mitochondrial inhibition reported here could contribute to clinical EPS. In SH-SY5Y cells and mouse neurons, respiration was significantly inhibited at concentrations of 5 μM and above. Although levels in patient serum are reported to be in the range of 0.1-1.3 μM (aripiprazole), 0.2-0.5 μM (brexpiprazole), and 0.09-0.4 μM (cariprazine) (37, 60–62), a pharmacokinetic study in mice showed that aripiprazole accumulated by approximately 9-fold greater levels in the brain (relative to serum levels) (63). Brain accumulation has also been reported for other antipsychotics, such as haloperidol (64). If this drug distribution behaviour translates into humans, partial agonists may concentrate at mitochondrial toxic levels and drive EPS. In mouse studies, the P-glycoprotein has been shown to regulate aripiprazole brain levels as it controls drug efflux; polymorphisms in this transporter may further sensitize particular patient groups to higher drug accumulation and toxicity (63, 65). Similarly, patients with pre-existing mitochondrial impairment (e.g. those with mitochondrial disease) are likely to be particularly susceptible to aripiprazole-induced mitochondrial toxicity (66). It would also be interesting to investigate whether specific mitochondrial DNA (mtDNA) haplotypes are correlated with an increased incidence of EPS following aripiprazole treatment: for example haplotypes conferring a predisposition to PD or reducing mitochondrial bioenergetic function (67–70)

We next considered how aripiprazole, brexipiprazole and cariprazine inhibit complex I. Recently IACS-2858, a structural analogue of the anti-cancer agent IACS-010759 currently in phase 1 clinical trials for acute myeloid leukaemia and solid tumours, was shown to inhibit complex I by binding tightly to a new inhibitory site near the entrance of the Q-site; thereby it inhibits ubiquinone reduction by a ‘cork in a bottle’ mechanism (71). Binding is mediated by a skeleton of aromatic/nonaromatic rings, with a similar architecture to the complex I inhibitors mubritinib and carboxyamidotriazole (72). Third-generation antipsychotic drugs also contain chains of aromatic and nonaromatic rings within their chemical backbone and therefore may also bind to the IACS-2858-site. Accordingly, we performed molecular docking analysis (Supp. 7). Interestingly, when aripiprazole, brexipiprazole and cariprazine were flexibly aligned to the fixed conformation observed in the IACS-bound mouse complex I cryo-EM structure (PDB: 7B93), they could be accommodated in the site, with protein-ligand interaction energies in line with equivalent values calculated here for IACS-2858 as well as mubritinib and carboxyamidotriazole (Supp. Fig. 7I). The same region of the site has also been observed, in different cryo-EM structures, to contain an adventitiously-bound detergent (73), a rotenone molecule and short-chain ubiquinone species (74), suggesting it is a good candidate for off-target drug binding. However, future structural studies are vital to unequivocally delineate how aripiprazole, brexipiprazole and cariprazine inhibit complex I. It will also be important to test substructures and closely related structural analogues to identify the chemistry conferring complex I binding; if such changes in chemistry are not essential for D2R-D3R binding, chemical modification may represent a possible means to dial-out off-target mitochondrial toxicity from these third-generation antipsychotics.

In conclusion, our results show that the third-generation antipsychotics aripiprazole, brexpiprazole and cariprazine induce off-target mitochondrial toxicity *via* selective inhibition of the Q-site of mitochondrial complex I. Whilst these drugs exhibit a more favourable side-effect profile relative to other antipsychotics, they still carry an unexpectedly high risk of EPS in adults and adolescents (75). We present novel and robust mechanistic toxicology data that may reconcile these clinical observations, given that complex I inhibition and mitochondrial dysfunction are strongly connected to motor dysfunction in neurodegenerative disease. To strengthen our understanding of the *in vitro*-to-clinic translation, further studies are essential that measure aripiprazole concentrations in the human brain (particularly in those patients experiencing debilitating EPS) and it will be of great interest to monitor the incidence of EPS if novel partial agonists are developed that lack complex I inhibition. These data will be critical to resolving the contribution of mitochondrial toxicity to clinical EPS and decreasing the incidence of side effects from antipsychotic medications.

## MATERIALS AND METHODS

### Materials

Aripiprazole, Clozapine, Quetiapine, Olanzapine, Paliperidone, Risperidone and Rotenone (Abcam), Dehydroaripiprazole (Santa Cruz Biotechnology), Didesmethylcariprazine (Clearsynth), Haloperidol, Iloperidone, Lurasidone, Zotepine (Sigma), Brexpiprazole and Cariprazine (Cayman Chemicals) were dissolved to desired stock concentrations in DMSO. Chlorpromazine (Sigma) was dissolved in distilled water, while Antimycin A, FCCP, Oligomycin A (Sigma) and Piericidin A (Cayman Chemicals) were dissolved in ethanol. All drug stock solutions were diluted into appropriate cell culture media to desired concentrations.

### Cell culture

The human neuroblastoma cell line SH-SY5Y (ATCC #CRL-2266) was maintained in Dulbecco’s Modified Eagle Medium/Ham’s Nutrient Mixture F12 (DMEM) (#31331028, Gibco), supplemented with 10% heat-inactivated Fetal Bovine Serum (FBS) and 1 mM sodium pyruvate (Gibco) at 37°C, 5% CO_2_ in a humidified incubator. Cells were routinely passaged twice weekly. Primary Human Cardiomyocytes (HCMs; PromoCell) were maintained in myocyte growth medium (PromoCell), supplemented with 11 mM glucose and supplement mix (5% FCS, 0.5 ng ml^-1^ epidermal growth factor, 2 ng ml^-1^ basic fibroblast growth factor and 5 μg ml^-1^ insulin). Cells were maintained at 37°C, 5% CO_2_ in a humidified incubator and passaged once per week. *Drosophila* Schneider 2 (S2) cells (Thermofisher) were maintained in Schneider’s *Drosophila* Medium (Gibco) supplemented with 10% FCS, at 26 °C in a non-CO_2_ incubator. Semi-adherent cells were passaged twice a week when approximately 85-90% confluence was reached.

### Glucose to galactose metabolic switching

SH-SY5Y cells were switched from standard growth medium to glucose-free DMEM (#11966025, Gibco) supplemented with 11 mM glucose or galactose, 10% FBS and 1 mM sodium pyruvate. Cells were grown in glucose or galactose-supplemented media for a period of 72 h prior to drug treatment.

### Isolation and culture of primary mouse neurons

Ventral midbrain and forebrain (76, 77) neurons were dissected from the brains of wild type (C57BL/6) day E13.5 mouse embryos, as previously described (76). Neurons were dissected in ice-cold DMEM + GlutaMAX (#10566016, Gibco) medium supplemented with 5% FBS. Following dissection, neurons were mechanically dissociated and then enzymatically digested using 1x TrypleExpress for 2 min at 37 °C. Dissociated midbrain and forebrain neurons were resuspended in neurobasal medium, prepared as previously described (Schmidt et al., 2012) and centrifuged at 200 x *g* for 5 min. Neurons were then seeded at the indicated densities into opaque 96-well tissue culture microplates (Corning™), Seahorse XF96 microplates (Agilent) or glass coverslips coated with Poly-D-lysine (50 μg ml^-1^, Sigma) in 24-well plates (Corning™). Neurons were maintained at 37 °C in a humidified 5% CO_2_ incubator for 7 d, with medium supplemented every 2-3 d.

### ATP assays

SH-SY5Y cells cultured in glucose or galactose-containing DMEM medium were seeded into opaque 96-well tissue culture microplates (Corning™) at a density of 2×10^4^ per well and incubated at 37 °C, 5% CO_2_ for 24 h. For experiments using primary neurons, mouse embryonic forebrain and ventral midbrain neurons were seeded at densities of 7.5×10^4^ and 1×10^5^ per well, respectively, and allowed to grow for 7 d at 37 °C, 5% CO_2._ On the day of the assay, media was aspirated from cells, and drugs added directly for a period of 18 h. Total cellular ATP was then measured using the Promega CellTiter-Glo® luminescent cell viability assay (#G7571), according to the manufacturer’s protocol. The average concentration of ATP (calculated from triplicates for each condition) was normalised to the average amount of ATP produced by DMSO-treated cells for each biological replicate. Data were fit to the standard dose-effect relationship (activity (%) = Bottom + (Top-Bottom)/(1+10^((LogIC50-X)*HillSlope)) using GraphPad Prism version 8.0.

### Cell proliferation assays

Cell proliferative capacity was measured in real-time using the xCELLigence® system (RTCA DP instrument; ACEA Biosciences), according to manufacturer’s instructions. SH-SY5Y cells were seeded into xCELLigence E-plates (ACEA Biosciences) at a density of 2.0×10^4^ cells/well (glucose-cultured cells) and 5.0×10^4^ cells/well (galactose-cultured cells) and incubated at 37 °C, 5% CO_2_, while impedance values (expressed as total cell index) were automatically monitored by the xCELLigence system. After an initial growth period of 24 h, impedance readings were paused to allow for addition of drugs to cells, and then resumed for a further 86 h. Average proliferative rates were calculated from the linear region of each growth curve, from triplicates per condition per biological replicate. These, along with final cell index values after the 110 h total growth period were normalised against those of DMSO-treated cells.

### Extracellular flux analyses

Measurements of real-time of oxygen consumption rates (OCR) were determined in whole cells using a Seahorse XF96 extracellular flux analyser (Agilent) as described previously (78), with the following modifications. SH-SY5Y cells were seeded into Seahorse XF96 microplates (Agilent) pre-coated with rat-tail collagen (30 µg ml^-1^, Sigma) at a density of 3.0×10^4^ cells/well in glucose-supplemented DMEM medium (containing 10% FBS, 1 mM sodium pyruvate). HCMs were seeded into uncoated XF96 microplates in glucose-supplemented myocyte growth medium at a density of 1.5×10^4^ per well. Mouse embryonic ventral midbrain and forebrain neurons were seeded into XF96 microplates pre-coated with Poly-D lysine, in neurobasal medium, at densities of 1.5×10^5^ cells/well and 1×10^5^ cells/well, respectively, and allowed to grow for 7 d. All cell types were incubated at 37 °C, 5% CO_2_ for 16 h after seeding. *Drosophila* S2 cells were seeded into uncoated XF96 microplates in Schneider’s medium at a density of 3.0×10^4^ per well, and incubated at 28 °C for 16 h without CO_2_.

On the day of the assay, growth medium was removed, and cells washed 3X in unbuffered serum-free DMEM Seahorse Assay Medium (32 mM NaCl, 2 mM GlutaMAX, 1 mM sodium pyruvate, 11 mM glucose). Cells were placed into a 37 °C non-CO_2__Jincubator for 1 h (apart from *Drosophila* S2 cells which were kept in a 28 °C non-CO_2__Jincubator). Antipsychotic drugs, diluted in Seahorse Assay medium to desired concentrations, were added directly, with cells then being incubated at 37 °C in a non-CO_2__Jincubator for 4 h. Following incubation, cells were placed into the Seahorse XF96 Analyser and OCR recorded, with canonical mitochondrial toxins injected sequentially at the indicated time- points. The canonical mitochondrial toxins oligomycin A (ATP synthase inhibitor), FCCP (mitochondrial uncoupler) and antimycin A with rotenone (complex III and I inhibitors, respectively) were loaded into sensor cartridge injection ports (Agilent) to yield final concentrations of 2 μM oligomycin A, rotenone, antimycin A and 500 nM FCCP for SH-SY5Y cells and 5 μM oligomycin A, 2 μM rotenone, antimycin A for mouse embryonic ventral midbrain and forebrain neurons. FCCP was added at 1 μM to forebrain neurons and 2 μM to ventral midbrain neurons. For HCMs, 2 μM oligomycin A, rotenone, antimycin A and 4 μM FCCP were used. Basal OCR was calculated by subtraction of antimycin A-independent OCR from the baseline OCR, prior to oligomycin A addition. Maximal OCR was calculated following by subtracting antimycin A-independent OCR from FCCP-stimulated OCR. Values were averaged from independent experiments (six technical replicates of each condition per experiment) and expressed as a percentage of those calculated for DMSO-treated cells. OCR measurements were made at 28 °C for *Drosophila* S2 cells, and 37 °C for all other cell types.

### Measurement of electron transport chain complex activity in permeabilised cells

To assess the activity of individual respiratory complexes in SH-SY5Y cells, cholesterol-dependent Permeabiliser XF Plasma Membrane Permeabiliser (PMP; Agilent) was used, essentially as described previously (79). To assess complex I activity, 10 mM pyruvate, 2.5_JmM malate and 1 mM ADP were injected into sensor cartridge ports, to stimulate complex I-linked respiration. This was followed by the injection of 2 μM piericidin A (to abolish complex I activity), followed by injection of_J10_JmM succinate and 1 mM ADP to stimulate complex II-linked respiration. All substrates were diluted in 1x Mitochondrial Assay Solution (MAS) buffer consisting of 70 mM sucrose, 220_JmM_Jmannitol, 10_JmM KH_2_PO4, 5_JmM MgCl_2_, 2_JmM_JHEPES, 1_JmM EGTA, at pH 7.4. Prior to assay, cells were washed twice in MAS buffer and drugs, diluted to desired concentrations in MAS buffer, immediately added to cells. PMP was then added directly at a final concentration of 1 nM in order to permeabilize the plasma membrane, whilst leaving mitochondrial membranes intact. OCR measurements were made using a Seahorse XF96 Analyser, with antimycin A-insensitive OCR subtracted from pyruvate/malate/ADP-stimulated OCR and succinate/ADP-stimulated OCR to give final OCR measurements for complex I and II-linked respiration. These values were then normalised to those obtained from DMSO-treated cells.

### Biochemical enzyme activity assays

All catalytic activity assays were conducted at 32 °C in 96-well plates using a Molecular Devices Spectramax 384 plus plate reader unless otherwise stated. Linear rates were measured for all assays, inhibitors added from DMSO stock solutions as required, and relevant inhibitor-insensitive rates measured using 1 µM piericidin A. Bovine and mouse membranes were prepared as described previously (80, 81), diluted to 20 µg mL^-1^ in 10 mM Tris-SO_4_ (pH 7.5) and 250 mM sucrose and supplemented with 3 µM horse heart cytochrome *c* (Sigma Aldrich). For measurement of NADH:O_2_ oxidoreduction, catalysis was initiated by addition of 200 µM NADH and monitored at 340 and 380 nm (ε_340-380_ = 4.81 mM^-1^ cm^-1^). Succinate oxidation was determined using a coupled enzyme assay (82) in the presence of 5 mM succinate, 60 µg mL^-1^ FumC, 300 µg mL^-1^ MaeB, 2 mM MgSO_4_, 1 mM K_2_SO_4_ and 2 mM NADP^+^. Bovine complex I was purified as described previously (83), and diluted to 0.5 µg mL^-1^ in 20 mM Tris-HCl (pH 7.5), 0.15% soybean asolectin (Avanti Polar Lipids) and 0.15% 3-[(3- Cholamidopropyl)dimethylammonio]-1-propanesulfonate (CHAPS, Merck Chemicals Ltd.). NADH:decylubiquinone (DQ), NADH:3-acetylpyridine adenine dinucleotide (APAD^+^), and NADH:ferricyanide (FeCN) oxidoreduction by complex I were measured using 200 µM NADH and 200 µM DQ, 100 µM NADH and 500 µM APAD^+^, or 100 µM NADH and 1 mM FeCN, respectively. Kinetic data were fit to the standard dose-effect relationship (activity (%) = 100 / (1 + (IC_50_) / ([inhibitor])^Hill^ ^slope^) using GraphPad Prism version 8.0, and IC_50_ values are reported with their standard errors. Sub-mitochondrial particles (SMPs) were prepared as described previously (81), and diluted to 10 µg mL^-1^ in 10 mM Tris-PO_4_ (pH 7.5), 250 mM sucrose and 2 mM MgCl_2_. ATP synthesis quantification was adapted from a previously described protocol (84). Briefly, the assay was performed at 20 °C using a Glomax 20/20 luminometer (Promega) by the luciferin/luciferase system supplemented with 25 µM P1,P5-di(adenosine-5’) pentaphosphate (AP5A; Sigma Aldrich), 3 µM IF1 (prepared as described previously (85), 100 µM ADP, and 20 µg mL^-1^ luciferase reagent (ATP Bioluminescence Assay Kit CLS-II, Roche). The reaction was initiated with 200 µM NADH, luminescence measured for 10 minutes, and the first derivatives calculated for the ATP synthesis rate.

### Cell death analyses

Cell death was assessed as described previously (86), with the following modifications. Glucose and galactose-cultured SH-SY5Y cells were seeded into 6-well plates (Corning™) at a density 7.0×10^5^ cells/well, incubated for 24 h at 37 °C, 5% CO_2_ and drugs then added at desired final concentrations, in glucose or galactose-medium. After 18 h treatment, cells were collected by trypsinisation and resuspended in glucose or galactose-medium, to allow recovery for 20 min at 37°C, 5% CO_2_. Cells were then diluted 1:5 in annexin binding buffer (1.8 mM CaCl_2_.2H_2_O, 1 mM MgCl_2_.6H_2_O, 5 mM KCl, 150 mM NaCl, 10 mM HEPES/NaOH, pH 7.4), followed by the addition of diluted Annexin V-FITC (synthesised in-house – (87)) and incubated for 30 min in the dark at room temperature. Samples were then placed on ice and 1 µl of the membrane impermeable, far-red fluorescent dye DRAQ7 (BioStatus) added to detect late-stage apoptotic or necrotic cells. Samples were analysed immediately using a BD FACSCanto™II (BD FACSDiva software) to quantify cell death.

### Western blotting

SH-SY5Y cells cultured in glucose-supplemented DMEM were seeded into 6-well tissue culture plates at a density of 7.5×10^5^ cells/well, while HCMs were seeded at a density of 1.3×10^5^ cells/well. Cell lysates were prepared in ice-cold RIPA buffer (Thermo Scientific™. #89901) supplemented with 100X Halt Protease and Phosphatase inhibitor cocktail for 15 min (Thermo Scientific™.#78440), and then centrifuged for 10 min at 13,000 x *g*, 4°C. Supernatant protein concentrations were normalised using a BCA protein assay (Pierce), according to the manufacturer’s protocol. Samples were prepared by diluting lysates in loading buffer containing 1% sodium dodecyl sulphate with β-mercaptoethanol, and heated for 5 min at 70 °C. These were run on 4-12% Criterion protein gels (Bio-Rad) at 100V for 1 h. Proteins were blotted onto nitrocellulose membranes using Trans-Blot^®^ Turbo™ Transfer System (Bio-Rad. #1704150). Membranes were blocked for 1 h at room temperature in buffer consisting of TBST (0.1% Tween-20, 10mM Tris-HCl, 150mM NaCl) and 5% dried skimmed milk powder (Marvel). Membranes were probed with primary antibodies against dopamine D2 receptor (Abcam, ab85367), dopamine D3 receptor (Abcam, ab155098), and vinculin (Abcam, ab130007), for 16 h at 4°C. This was followed by a 1 h incubation at room temperature with anti-mouse or anti-rabbit secondary antibodies conjugated with horseradish peroxidase (HRP; Sigma, A8924). Blots were developed using a Konica SRX101A X-Ray Film Processor.

### Immunofluorescence

For analysis of TH expression in the ventral midbrain and forebrain, whole brains from E13.5 embryos were cryosectioned on sagittal planes, fixed with 2% PFA for 30 min at room temperature, and then simultaneously permeabilised and blocked for 30 min at room temperature with a buffer containing 10% donkey serum (Jackson ImmunoResearch, UK) and 0.1% Triton X-100 (Sigma). Rabbit anti-tyrosine hydroxylase (TH) was added overnight in blocking solution at 4°C to whole brain cryosections. The next day, neurons were washed in PBS and incubated in the dark for 1 h with donkey anti-rabbit Alexa Fluor 555 (1:500). Neurons were washed 3 times in PBS then stained with Hoechst to identify nuclei. Stained cells were imaged using a confocal microscope (LSM 880, Zeiss).

### Drosophila strains

Fly stocks were maintained on standard cornmeal agar media at 25°C. The strain used was *w^1118^.* All behavioural experiments on adult flies were performed using males.

### Isolation of fly mitochondria

Mitochondria were isolated from freshly eclosed adult *Drosophila,* according to the manufacturer’s protocol (Sigma). Briefly, 50 flies (male and female) were placed onto ice and gently homogenized using a Dounce homogenizer, with 10 strokes per extraction. This was followed by a 5 min centrifugation of supernatants at 600 x *g* and 10 min centrifugation at 11000 x *g*, 4°C. Pellets were resuspended in complex I assay solution (1X MAS buffer supplemented with 10 mM pyruvate and 5 mM malate) at 4°C, and total protein content normalized using a Bradford assay.

### Behavioural analyses in *Drosophila*

Freshly eclosed male flies were maintained on standard cornmeal agar media supplemented with 1 mM aripiprazole, 1 mM rotenone or 0.5% (v/v) DMSO for 5, 14, 21 or 28 days. For the climbing assay, 20 male flies were placed into glass columns (23 cm long, 2.5 cm in diameter) that were lined with nylon mesh (250 micrometres, Dutscher Scientific) and marked with a line at 15 cm. After a 30–60 min recovery from CO_2_ anaesthesia, flies were gently tapped to the bottom of the vial, and the time required for 5 flies to climb above the marked line was recorded. For each experiment, at least three cohorts of 20 flies from each treatment group were scored (with three individual biological repeats for each group). Climbing latency was defined as the average time in seconds for 5 flies to climb above the marked line, with averages being pooled together from each group of 20 per treatment condition. The climbing latencies for rotenone and aripiprazole-fed flies were then normalised and expressed as a fold-change relative to that obtained for DMSO-fed flies (which was set to 1.0). For the activity assay, individual males were loaded into Drosophila Activity Monitors (DAM5) within 8 × 65-mm glass Pyrex tubes (Trikinetics, Waltham, MA, USA) containing fly food supplemented with 1 mM aripiprazole or 0.5% (v/v) DMSO. The flies were maintained at 25 °C under a 12-hour light:12-hour dark (LD) cycle. Activity data were analysed using the Sleep and Circadian Analysis MATLAB Program (SCAMP) developed by the Griffith lab (88). The analyses were performed for 7, 14 and 21 days starting at the first ZT0 to allow acclimation. At least 16 flies of each treatment were used.

### Transmission electron microscopy

For transmission electron microscopy, adult fly brains or thoraces were fixed for 2 h in 0.1 M sodium cacodylate buffer (pH 7.4) containing 2% paraformaldehyde and 2.5% glutaraldehyde (at room temperature). Then, samples were post-fixed for 1 h in a solution containing 0.25% osmium tetroxide/0.25% potassium ferrocyanide and 1% tannic acid. After fixation, the samples were stained *en bloc* with 5% aqueous uranyl acetate overnight at room temperature; then, dehydrated via a series of ethanol washes and embedded in TAAB epoxy resin (TAAB Laboratories Equipment Ltd., Aldermas-ton, UK). After polymerization at 65°C for a few days, the ultrathin-sections (approximately 60 nm) obtained by Ultramicrotome (Leica Ultracut UCT, Vienna Austria) were mounted in EM grids, stained with lead citrate, and then observed by FEI Talos F200C 200kV transmission electron microscope (Thermo Fischer Scientific, Oregon USA) with Ceta-16M CMOS-based camera (4kx4k pixels under 16bit dynamic range). To investigate mitochondrial ultrastructural changes in flies fed DMSO, rotenone or aripiprazole, individual mitochondria were counted and ranked as either healthy, partially damaged or very damaged, and then normalised against the total number of mitochondria in that field of view. Partially damaged mitochondria were quantified as those exhibiting small areas of ultrastructural disruption, whereas very damaged mitochondria displayed areas of widespread cristae fragmentation. Images from each respective group were counted by one individual and blinded to eliminate the risk of sample bias.

### Small-molecule conformational analyses

A selection of noncanonical complex I inhibitors including aripiprazole were flexibly aligned to the fixed conformation of complex I structure (PDB entry 7B93) using AutoDock Vina (89). Charges were added to the ligands and receptors using AutoDockTools (https://www.ncbi.nlm.nih.gov/pmc/articles/PMC2760638). Simulations were performed at an exhaustiveness value of 300 on a high-performance computer with 64 central processing units. The top 10 binding poses were analysed, and the best conformer was selected by comparing the protein-ligand interaction energies calculated. The detailed workflow for the docking analysis, as well as the PDB coordinates of the results are available in GitHub (m1gus.github.io/Aripiprazole/).

### Statistical analyses

Statistical analyses were performed using GraphPad Prism 8.0 (www.graphpad.com). The data are presented as the mean values, and the error bars indicate ± SD or SEM. In violin plots, the solid line represents the median, whereas dotted lines represent the quartiles. The number of biological replicates per experimental variable (n) is indicated in either the respective figure or figure legend. Significance is indicated as * for p < 0.05, ** for p < 0.01, *** for p < 0.001, and **** for p < 0.0001.

## Acknowledgements

We would like to thank Dr Lucia Pinon (MRC Toxicology Unit) for assistance with flow cytometry and confocal microscopy, Tim Smith and Maria Guerra Martin for help with processing of Electron Microscopy samples and Tim Ashby and Munisha Patel for preparing the fly food.

## Data availability

All datasets are securely archived and are available to researchers on request.

