## Supplemental Figures S1-S7 for "The antipsychotic medications aripiprazole, brexpiprazole and cariprazine are off-target respiratory chain complex I inhibitors"

**This PDF file includes:**

Figures S1 to S7

### Supplementary Figure 1

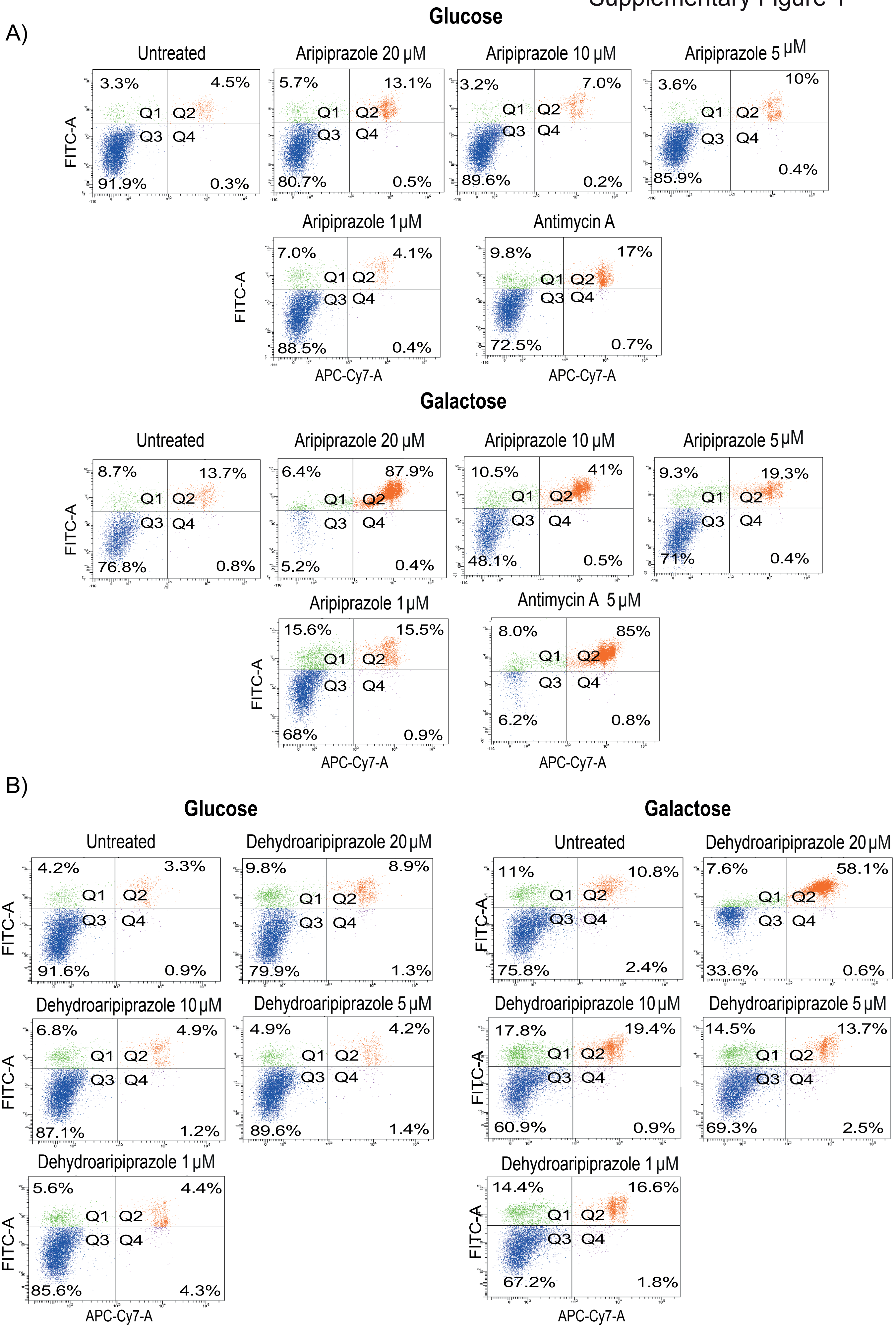

**Supplementary Figure 1. Aripiprazole and dehydroaripiprazole induce cell death in galactose-conditioned SH-SY5Y cells**

Cell death quadrants obtained from glucose or galactose-conditioned SH-SY5Y cells treated for 18 h with increasing concentrations of (A) aripiprazole or (B) dehydroaripiprazole. Cells were stained with annexin-V and DRAQ-7 and analysed by flow cytometry to identify early-stage apoptotic (green) or late-stage apoptotic (red) cells.

#### Supplementary Figure 2

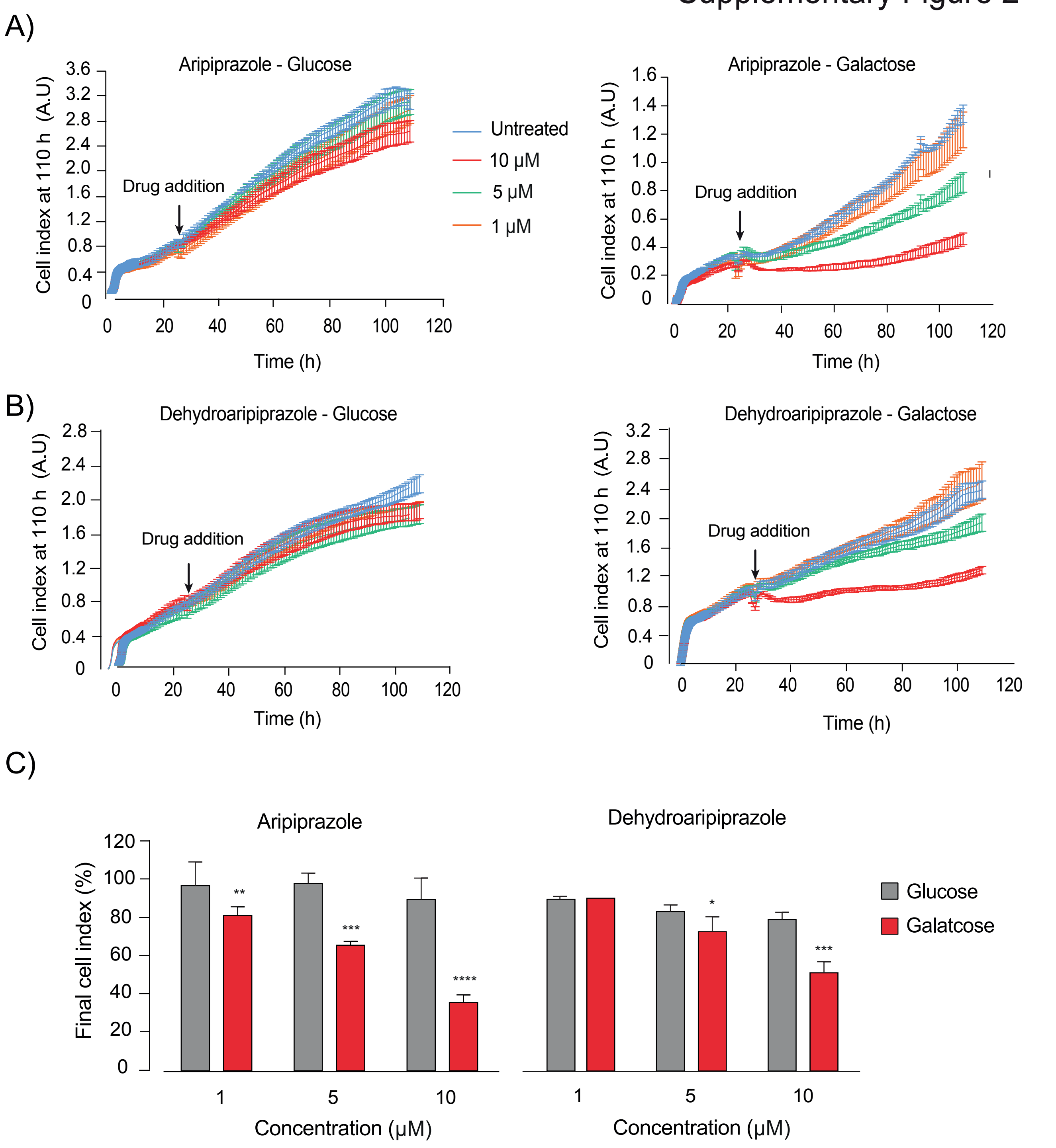


**Supplementary Figure 2. Aripiprazole and dehydroaripiprazole reduce the proliferation of galactose-conditioned SHSY5Y cells**
Representative proliferation traces for glucose or galactose-conditioned SH-SY5Y cells treated for 72 h with (A) aripiprazole or (B) dehydroaripiprazole at indicated concentrations. The point of drug addition is indicated with an arrow. (C) Quantification of final cell index values following 86 h exposure of glucose and galactose-conditioned SH-SY5Y cells to aripiprazole or dehydroaripiprazole (mean ± SEM from 3 independent experiments, asterisks, one-way ANOVA with Dunnett's multiple comparison test, normalised to control).

### Supplementary Figure 3

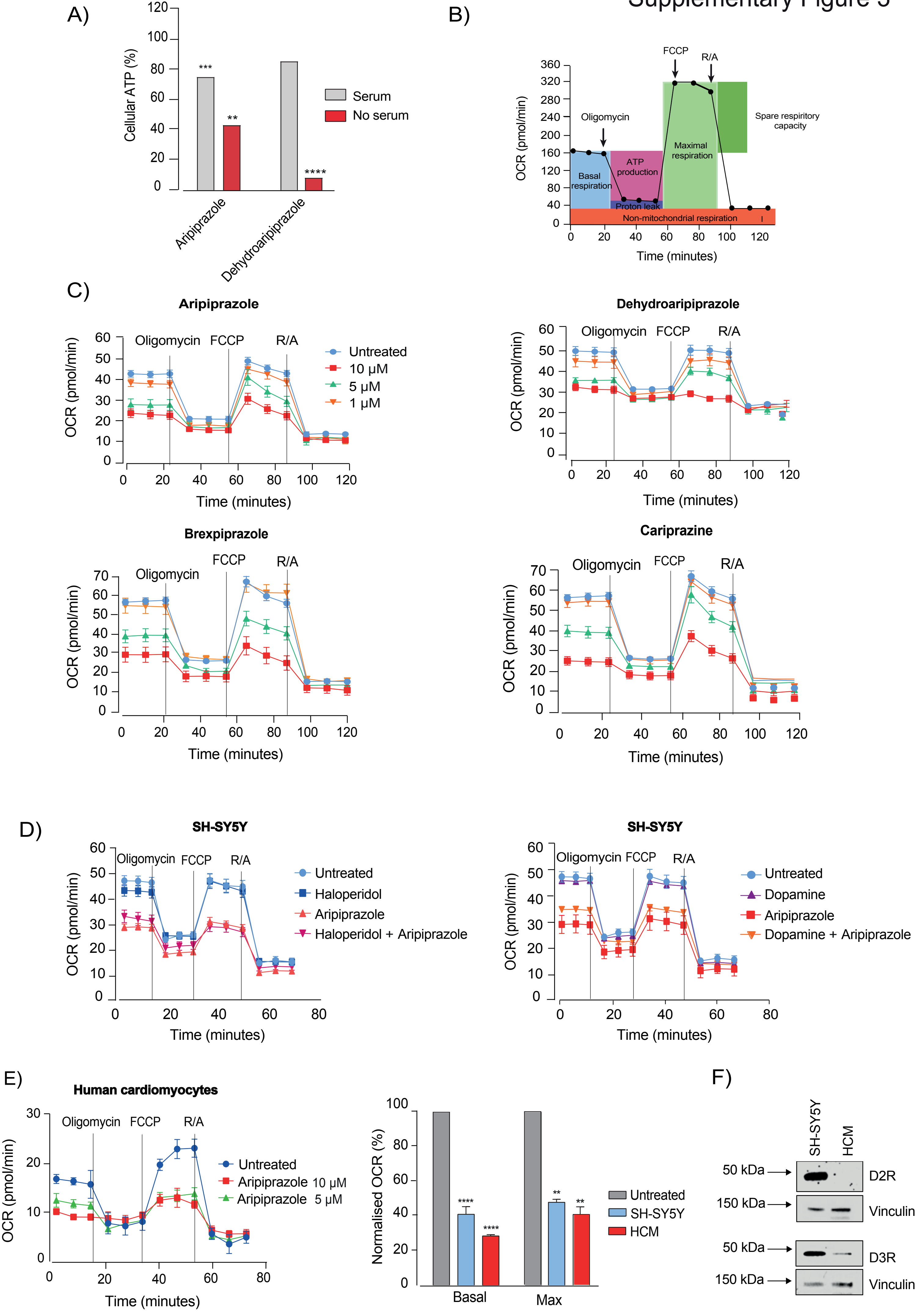

**Supplementary Figure 3. Aripiprazole and structurally related antipsychotic drugs are direct mitochondrial toxins**

(A) Normalised cellular ATP measurements from galactose-conditioned SH-SY5Y cells cultured in media supplemented with 10% FBS or no FBS. Cells were exposed to 10 μM aripiprazole or dehydroaripiprazole for 4 h (mean ± SEM from 3 independent experiments, asterisks, one-way ANOVA with Dunnett's multiple comparison test, normalised to control). (B) Representative Seahorse trace for a standard mitochondrial stress test indicating key bioenergetic parameters. (C) Representative Seahorse traces from SH-SY5Y cells treated for 4 h with the indicated antipsychotic drugs. Each OCR measurement is presented as a mean ± SEM from 6 independent wells per treatment. (D) Representative Seahorse traces for SH-SY5Y cells treated with 50 μM dopamine or haloperidol, or 10 μM aripiprazole. Cells were pre-treated for 20 minutes with dopamine or haloperidol prior to aripiprazole addition. (E) Representative Seahorse trace for HCMs treated for 4 h with 5 or 10 μM aripiprazole. Quantification of basal and maximal OCR in HCMs compared to SH-SY5Y cells after 4 h treatment with 10 μM aripiprazole (mean ± SEM from 2 independent experiments, asterisks, one-way ANOVA with Dunnett's multiple comparison test, normalised to control). (F) Western blots showing expression of D2R/D3R in HCMs compared to SH-SY5Y cells. Vinculin was used as a loading control.

Supplementary Figure 4

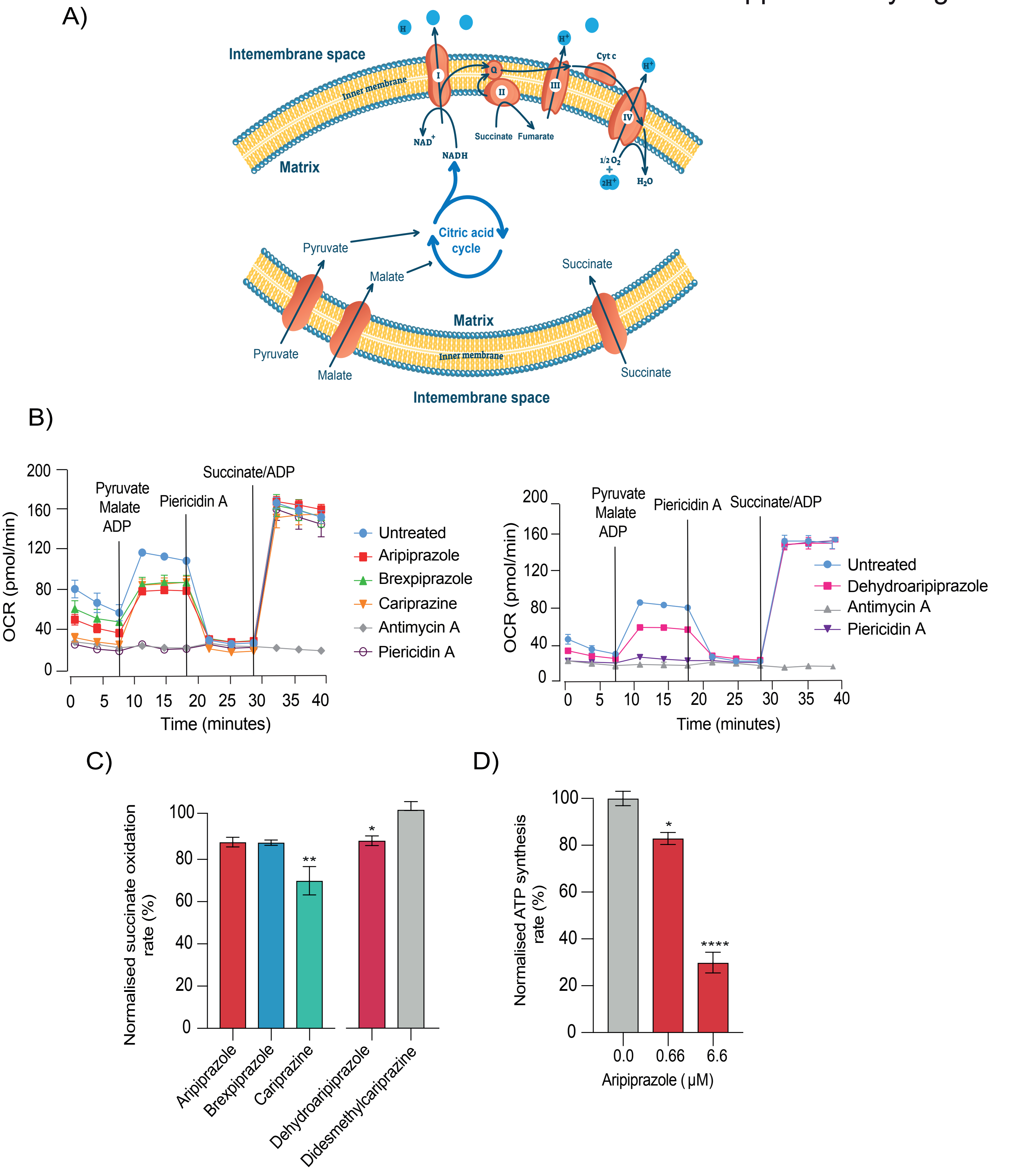


**Supplementary Figure 4. Aripiprazole and structurally-related antipsychotic drugs are mitochondrial complex I Q-site inhibitors**

(A) Schematic of the reactions that occur in assays on permeabilized cells used to dissect the activity of respiratory-chain complexes. Exogenous pyruvate and malate are metabolised by the TCA cycle, generating NADH to stimulate complex I-dependent respiration. Exogenous succinate directly stimulates complex II-dependent respiration. (B) Representative Seahorse trace obtained from SH-SY5Y cells treated with 10 μM of indicated antipsychotic drugs and 5 μM piericidin or antimycin A. Each OCR measurement is presented as a mean ± SEM from 6 independent wells per treatment. (C) Succinate:O2 oxidoreduction rates from bovine mitochondrial membranes exposed to 100 μM aripiprazole, brexpiprazole, or cariprazine, or 50 μM dehydroaripiprazole or didesmethylcariprazine (mean ± SEM from 3 independent wells per treatment, asterisks, one-way ANOVA with Dunnett's multiple comparison test, normalised to control). (D) ATP synthesis measurements from bovine sub-mitochondrial particles exposed to the indicated concentrations of aripiprazole (mean ± SEM from 3 inde-pendent experiments per treatment, asterisks, one-way ANOVA with Dunnett's multiple comparison test, normalised to control).

A)

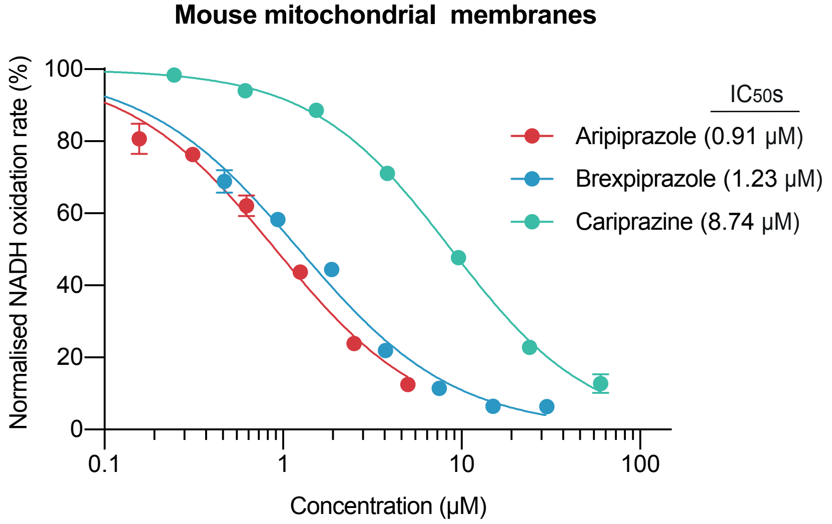


B)

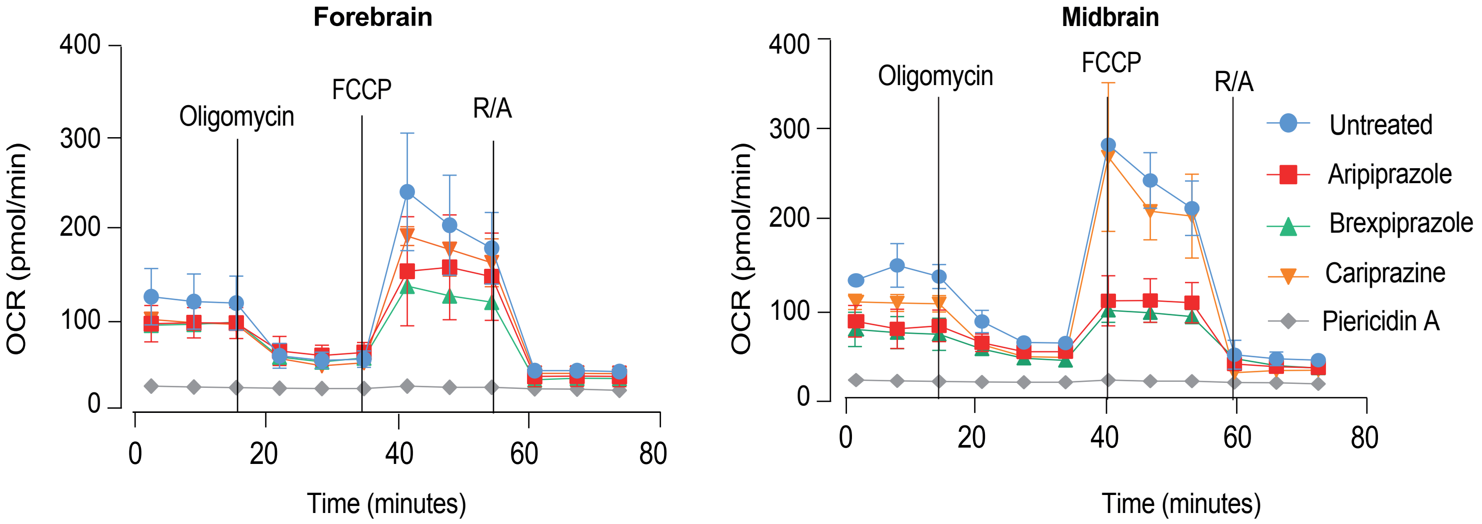


**Supplementary Figure 5. Aripiprazole and structurally related antipsychotic drugs are mitochondrial toxic in primary mouse neurons**

(A) Normalised NADH:O2 oxidoreduction rates of mouse mitochondrial membranes exposed to varying concentrations of aripiprazole, brexpiprazole and cariprazine. Respective IC50 values are indicated in brackets. (B) Representative OCR profiles from mouse ventral midbrain and forebrain neurons (e13.5) cultured for 7 d. Neurons were treated with aripiprazole, brexpiprazole, cariprazine, or piericidin A (5 µM) for 4 h. Each measurement is presented as a mean ± SEM from 3 independent wells per treatment (ventral midbrain) and 5 independent wells per treatment (forebrain).

### Supplementary Figure 6

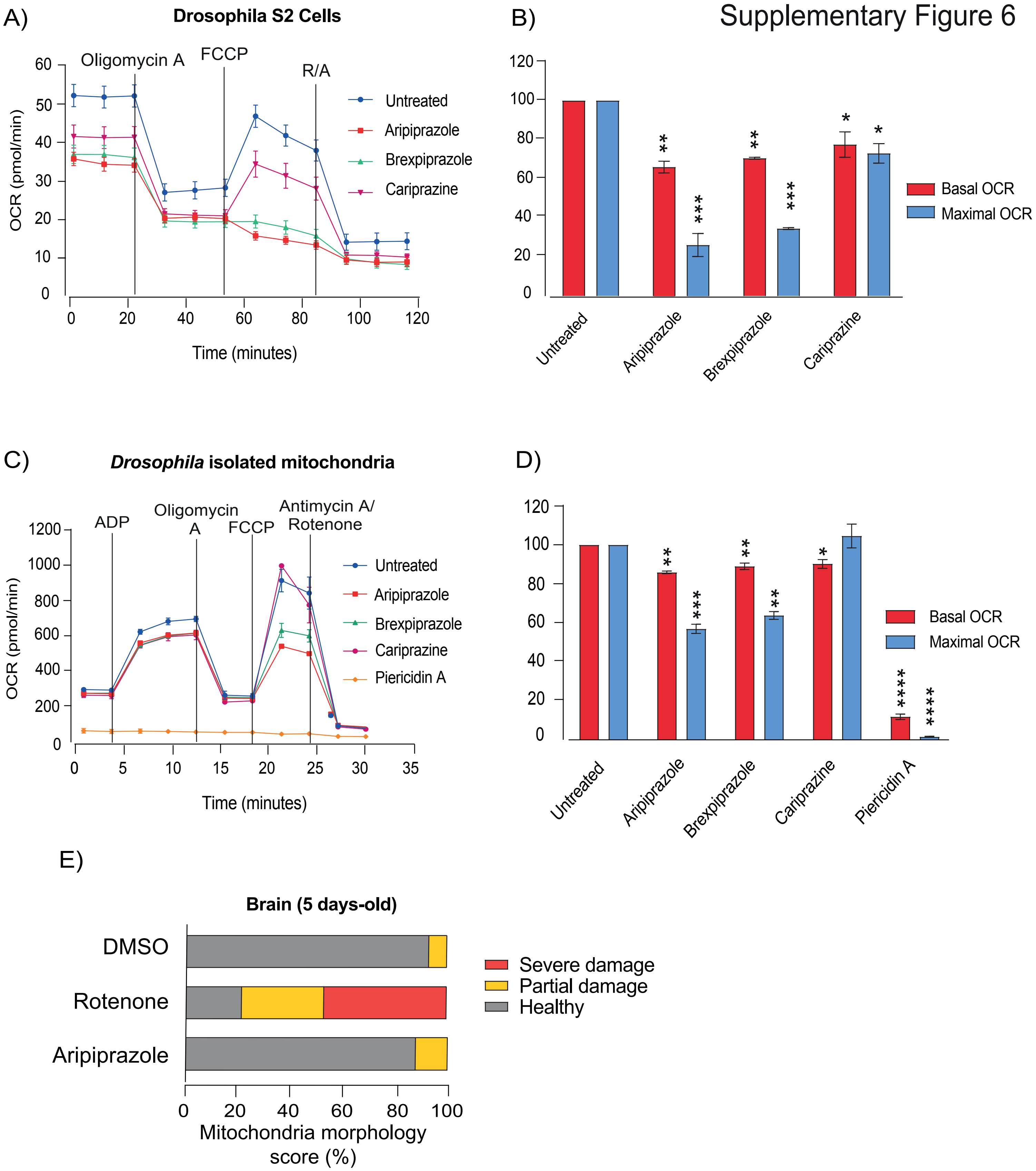


**Supplementary Figure 6. Aripiprazole displays mitochondrial toxicity in Drosophila melanogaster**
(A) Representative Seahorse trace from Drosophila S2 cells treated with 10 μM of indicated drugs for 4 h. Each measurement is presented as a mean ± SEM from 6 independent wells per treatment. (B) Corresponding normalised basal and maximal OCR measurements (mean ± SEM from 3 independent experiments, asterisks, one-way ANOVA with Dunnett's multiple comparison test, normalised to control). (C) Representative Seahorse trace from isolated Drosophila mitochondria exposed to 10 μM of indicated drugs. Respiration was measured immediately after mitochondria had been exposed to drugs. (D) Corresponding normalised basal and maximal OCR measurements (the data are mean averages ± range from 2 independent experiments with 2 technical repeats, asterisks show the results from one-way ANOVA with Dunnett's multiple comparison test, normalised to control). (E) Quantification of mitochondrial damage in Drosophila melanogaster brain. Adult flies were treated with DMSO (0.5%, v/v) aripiprazole or rotenone (both 1 mM) for 5 d.

### Supplementary Figure 7

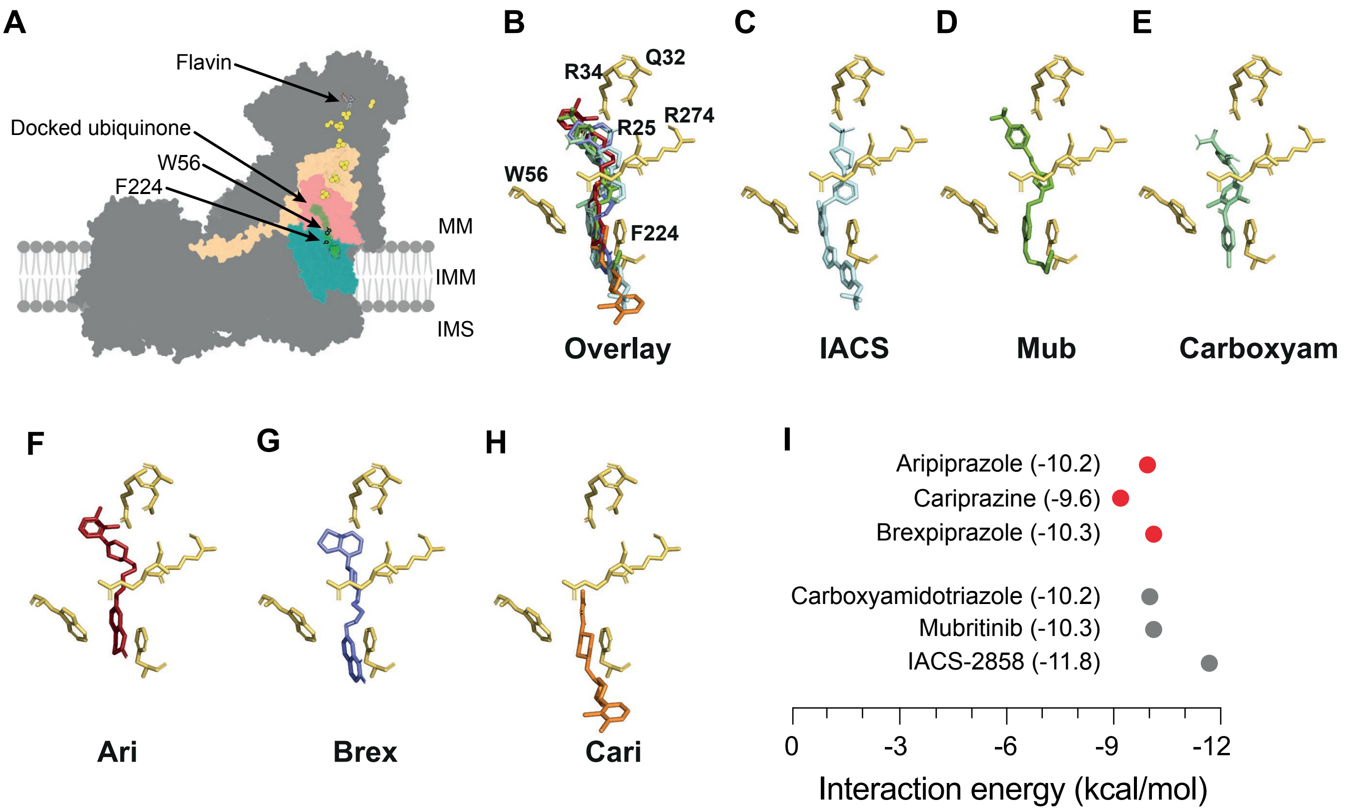


**Supplementary Figure 7. Molecular docking analysis of aripiprazole, brexpiprazole and cariprazine at the IACS-2858 binding site of complex I**

(A) A surface representation of mouse complex I (PDB:6ZR2). Blue mesh, docked ubiquinone-10; yellow/orange spheres, iron-sulphur clusters; MM, mitochondrial matrix; IMM, inner membrane; IMS, inter-membrane space. NDUFS2, NDUFS7, ND1 and a docked ubiquinone are shown in beige, salmon, teal and green, respectively. (B-H) Molecular docking simulating the alignment of all ligands overlaid (B), IACS-2858 (C), mubritinib (D), carboxyamidotriazole (E), aripiprazole (F), brexpiprazole (G), and cariprazine (H), with the residues involved in the binding of IACS-2858 as described in (71), at the entrance of the ubiquinone-binding site. (I) Protein-ligand interaction energies were calculated using Autodock Vina (see Methods).
